# Heatwaves and turbidity influxes have interacting and lasting effects on the foraging behaviour of three-spined sticklebacks *Gasterosteus aculeatus*

**DOI:** 10.64898/2026.10.15.759403

**Authors:** Cait Newport, Ellen Morley, Richard Creswell, Michelle Jackson, Ben Lambert

## Abstract

Freshwater ecosystems support a disproportionately high share of fish species and biodiversity yet are among the most threatened ecosystems globally and are increasingly being exposed to multiple, co-occurring environmental stressors. In addition, predicting the combined impacts of environmental stressors remains a major challenge, particularly when stressors differ in how quickly they arise and how long their effects persist. Understanding how these stressors interact across time is critical for predicting their ecological consequences. Here, we examine how warming and turbidity—two widespread and intensifying stressors—interact to influence predator–prey dynamics in the three-spined stickleback (*Gasterosteus aculeatus*). Using a fully factorial experiment, fish were exposed to ambient, warmed, turbid, or combined conditions for short (one week) or prolonged (six weeks) periods, before feeding rates were quantified across prey densities using a novel stochastic Bayesian model. We found that foraging behaviour was shaped primarily by prior exposure rather than the immediate experimental conditions. Short-term exposure to both stressors increased feeding rates, whereas prolonged warming reduced them. In contrast, turbidity continued to increase feeding rates over both time periods and, when combined with warming, offset the negative effects of extended heat exposure. Changes in feeding performance at the individual level may propagate through food webs, potentially triggering trophic cascades that reduce biodiversity. Given the exceptional biodiversity of freshwater systems, such dynamics are of particular concern. We also found that behavioural changes persisted even after stressors were removed, indicating that exposure history can have lasting effects. These findings suggest that short-term experiments, such as the functional response, may actually be measuring subject experiential history rather than responses to specific experimental conditions.

## 1 Introduction

Freshwater ecosystems support a disproportionate share of global biodiversity [1] and are among the most threatened environments on Earth [2, 3]. Their close proximity to human activity combined with limited options for organisms to relocate, can result in them being exposed to multiple, often simultaneous, environmental stressors. Among the most pervasive of these stressors are rising temperatures and increasing turbidity. Climate warming is intensifying not only average freshwater temperatures [4, 5], but also the frequency and duration of heatwaves [6]. Average lake heatwave temperatures are predicted to increase from 3.7°C to 5.4°C and the duration of heatwaves is expected to rise from an average of one week to 12 weeks, under a high-greenhouse-gas-emission scenario [6]. Simultaneously, concentrations of both organic and inorganic turbidity are increasing in many freshwater ecosystems. This increase is driven by both direct human activities, including industrial and agricultural runoff leading to eutrophication [7], dam discharges, and dredging [8, 9], and by climate change, which is intensifying extreme precipitation, flooding events, and forest fires [10, 11]. As rates of both warming and turbidity continue to increase globally, these stressors are more likely to co-occur [12].

In fish, the physiological effects of warming have been the focus of many studies, and warming has been shown to elevate metabolic rates and oxygen demand, while simultaneously reducing the oxygen carrying capacity of water [13, 14, 15]. Turbidity, on the other hand, can affect physiology but also sensory function, with impacts that are species, turbidity source, and ecological context dependent. For example, turbidity can damage gills [16], reduce visual cue reliability through light scattering [17], and interfere with olfactory cue detection [18]. As a result, turbidity can alter how fish acquire information about their environment, including cues related to prey capture and mate selection [19].

Environmental stressors vary not only in intensity but also in the timescales over which they act. In freshwater systems, turbidity can increase rapidly (e.g. minutes to hours) whereas warming develops comparatively slower due to the heat capacity of water (hours to days). Both stressors can persist for extended periods under conditions such as eutrophication or prolonged heatwaves. These differences in temporal dynamics mean that fish may experience stressors either as short-term pulses or prolonged exposures, with potentially different consequences for behaviour, and ultimately survival [but see 20]. Importantly, prior exposure to environmental stress may modify subsequent responses, such that changes in behaviour may reflect not only current conditions but also exposure history [21, 22, 23].

Alterations in individual behaviour are particularly consequential for trophic interactions, where changes in individual behaviour can scale up to influence population and community dynamics. Food webs are largely understudied in a multiple stressor context (but see [24], yet they are important to understanding how individual responses cascade to influence the other species they interact with, either as predator or prey. One way to quantify such effects is through functional response experiments, which describe how prey consumption varies with prey density and allow estimation of key behavioural parameters such as predator attack rate and handling time [25, 26]. This framework provides a mechanistic link between environmental conditions and predator–prey interactions, making it well suited to assessing how multiple stressors alter foraging and its potential long-term effects on food webs and ecosystem dynamics.

We used the three-spined stickleback (*Gasterosteus aculeatus*), a visually oriented mesopredator, and a well-established model species in behavioural and ecological research. Sticklebacks are widely distributed across freshwater systems in the Northern Hemisphere [27] and have been extensively used to study foraging, sensory ecology, and responses to anthropogenic stressors [e.g. 28, 29, 30, 31]. This species is known to have a wide thermal tolerance but the degree of warming predicted under a high-greenhouse-gas-emission scenario for an extended period still causes higher mortality and higher energy requirements [32].

Three-spined sticklebacks are also sensitive to turbidity. This species relies heavily on visual cues for a range of behaviour including predation and sexual selection [29, 33] and turbidity has been shown to disrupt these behaviours. For example, turbidity can weaken sexual selection [34], alter courtship displays [35], and affect shoaling [36]. It also influences foraging. Moderate levels of bentonite clay induced turbidity can reduce reactive distance to prey, but without necessarily decreasing overall consumption [37], and in some cases increasing attack rates [38]. The lack of effect on consumption may reflect a shift towards the use of other sensory modalities, such as olfaction, which can become more important when visibility is reduced [39]. However, algal-induced turbidity has been shown to reduce both prey consumption and prey selectivity in this species [40]. Therefore, the responses of three-spined stickleback to warming and turbidity are complex and may depend on a range of factors including exposure history, magnitude of the stressor, turbidity type, and the mechanism underlying behavioural change [41]. It remains unclear how these stressors will interact: effects may be additive through shared physiological pathways, or turbidity-induced sensory impairment may be mitigated by the use of alternative sensory systems.

To disentangle the effects of stressor type and exposure duration, we experimentally tested how warming and turbidity, individually and in combination, affect fish foraging behaviour. In this fully factorial experiment, fish were exposed to ambient, warmed, turbid, or combined warming and turbid treatments, for either one week, or six weeks. Following these exposure periods, we quantified prey consumption using a functional response approach, measuring feeding rates across seven prey densities during a standardized one-hour feeding trial conducted under each of the four experimental conditions. This design allows us to assess whether behavioural responses are driven primarily by immediate environmental conditions or by the duration of prior exposure.

## 2 Methods

### 2.1 Study species

A total of 672 wild-caught sticklebacks from several freshwater waterways within England were used. Authors CN and EM caught 158 fish from Lincolnshire, UK (November 2021). All remaining fish were purchased from a local supplier (DC Freshwater Fish Ltd.) in Nottingham. All experimental procedures and animal husbandry protocols were reviewed and approved by the University of Oxford, Department of Biology Local Ethical Review Committee (AWERB: APA/1/5/ ZOO/NASPA/Newport&Jackson/Sticklebacks) prior to the commencement of this study. Fish sex was not identified as the study occurred outside of the UK breeding season so external indicators were not visible. Experiments were undertaken between November 2021 and March 2022.

### 2.2 Acclimation

During acclimation, fish were held in a flow-through aquarium system. Each tank was 58L (350 × 300 × 600 mm) and fish were maintained at a stocking density of <0.3 fish/litre (15 - 17 individuals per tank) with a 12:12 hr photoperiod. Water chemistry was regularly measured and maintained at healthy levels (salinity: 0.002-0.004; pH: 7-7.8; Ammonia: 0ppm; Nitrite: 0ppm; Nitrate: >10ppm). All tanks had at least one plastic plant to provide shelter.

Fish were acclimated within the flow-through aquarium system to one of four conditions (Fig. 1): 1) Ambient: 15°C clear water, 2) Warmed: 20°C clear water, 3) Turbid: 15°C turbid water, and 4) Mixed: 20°C turbid water. Turbid water was created by adding 0.47g of bentonite clay per litre, generating a turbidity level of 28.94±0.91 Nephelometric Turbidity Units (NTU). Bentonite clay was selected as a turbidity source as it appears similar in colour and particle size to the turbidity source found at our field site, and it has been used in previous studies with this species [37]. During acclimatisation periods, turbidity was checked twice daily and resuspended by manual mixing with a hand. Temperature was controlled using a water chiller for cold water or ambient room temperature for warm water. Temperature was recorded throughout using HOBO loggers (HOBO UA-001-64). For each condition, fish were assigned to either a one- or six-week acclimation period. Fish were fed twice daily *ad libitum* with defrosted bloodworms and brine shrimp (ratio 2:1).

**FIGURE 1.**
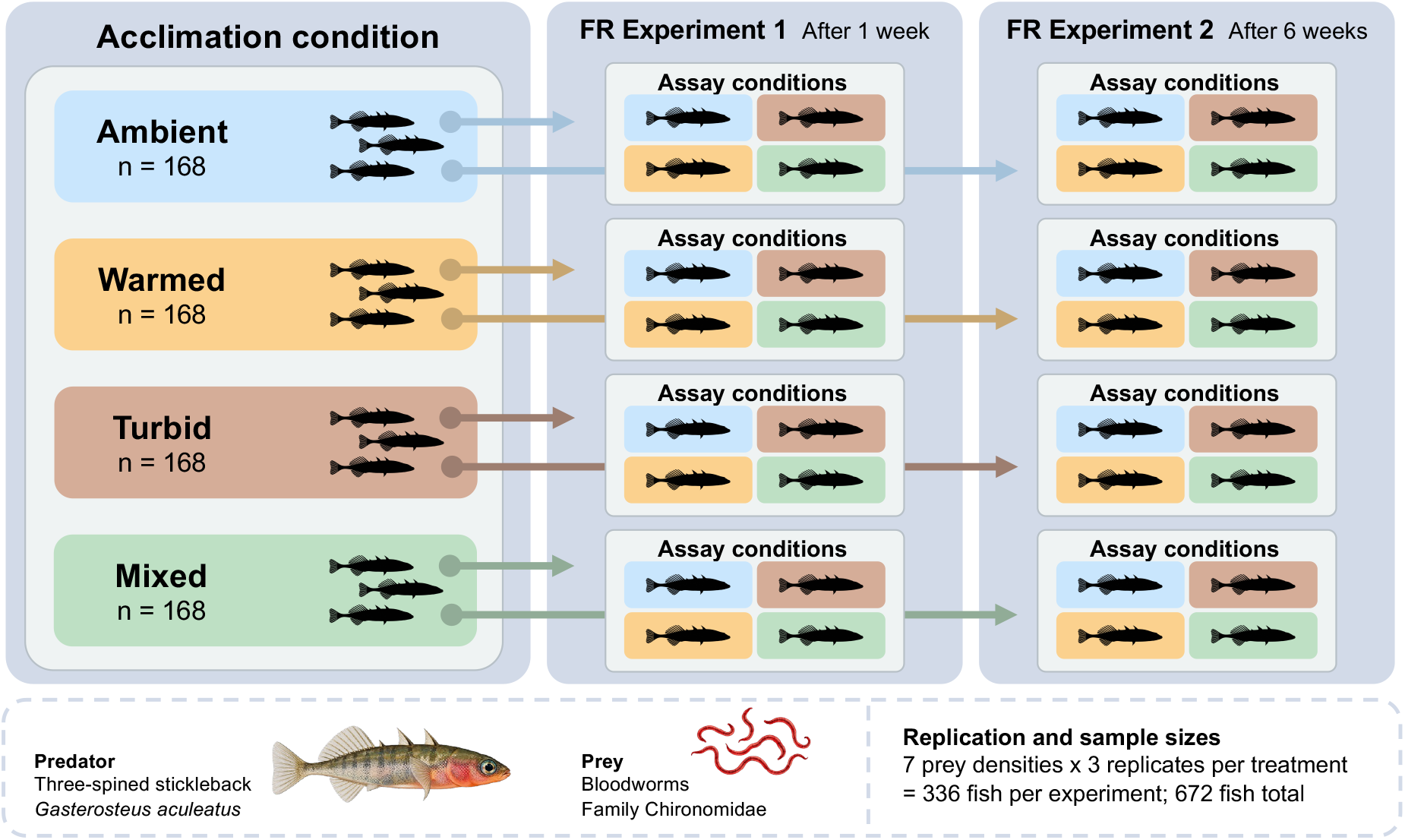
Illustration of the fully factorial experimental sampling procedure. FR refers to the Functional Response experiments. Illustrations of fish and bloodworms have been made using chatGPT (5.6)

### 2.3 Functional response (FR) experiment

Each experimental tank contained one of seven possible prey densities (5,10,15, 20, 25, 30, 40 live bloodworms) and one of four water treatments (ambient, warmed, turbid, mixed). The warming treatment was achieved by placing the tank in either a 15° C or 20° C water bath. An extra tank in each water bath contained a submerged logger, measuring temperature at 15-minute intervals, and was used to verify temperature consistency. The experimental turbidity level was 99.28±0.328 NTU (mean ± sd), generated using 1.02 g/L of bentonite clay. Note that the turbidity level was lower during acclimation to ensure fish could be observed during twice daily animal health checks. The experimental turbidity level of ~100 NTU is similar to the level used in previous studies [e.g. 42,43] and maximal levels measured in potential *G. aculeatus* habitats in the United Kingdom [44]. To reduce the effect of particle settlement, turbidity was resuspended in all tanks using an electric frother midway through the one-hour experimental period. To ensure all fish experienced the same treatment, even tanks with clear water were mixed. At the end of each experiment, turbidity was remeasured in each turbid tank (mean turbidity = 42.037±1.6 NTU).

Fish were not fed within 24 hours before experiment began. Experiments begun by transferring fish from their acclimation tanks to individual experimental tanks (250×158×128mm). Fish were caught by hand net and placed in a clear bag with 200mL of water from their home tank. The bag was then suspended for 15 minutes within a 3L water bath matching the water temperature that the fish were being moved to. This step was introduced to reduce the risk of stress caused by large temperature changes. After this stage, fish were then transferred directly into their experimental tank by hand net. Fish were given one hour to eat freely and at the end of this period the number of bloodworms remaining in the tank was counted. Three fish replicates were conducted for each treatment condition and prey density. Because of the time required to run experiments, replicates were conducted on consecutive days (28 fish run per day). After 1 hour, fish were removed from their tank by hand net and transferred directly to a solution of Tricaine mesylate (MS222; 1g MS222 + 3g bicarbonate soda per litre of water) and euthanised and the remaining prey items were counted. Fish total length, standard length and weight were measured.

We recorded trials using a GoPro Hero5 mounted on a second opaque tank placed directly on top of the tank containing a fish. Only a segment of each trial was filmed due to limitations in camera availability (7 cameras used). Each fish on a given experimental day was randomly assigned to one of four filming groups and each group was filmed for 8 minutes of the experiment (1080 × 1920 pixels; 60 frames per second, fps). These videos were not included in the experimental analysis but the procedure is reported because the presence of recording equipment could influence fish behaviour.

### 2.4 Statistical methods

#### 2.4.1 Mechanistic model

In order to represent the feeding behaviour of fish in our experiments, we developed a novel mechanistic model. Our model is inspired by the functional response literature, where the original form of the mathematical models were devised by Holling in 1959, and were justified through simple laboratory-based experiments that timed how long it took human participants to pick up “discs” from a table [25]. The mathematical forms, and expanded versions of these, have been used as part of differential equation-based descriptions of predator-prey systems [45], which has underpinned popular software used to estimate functional responses from data [e.g. 46]. Existing approaches, to our knowledge, however, do not explicitly allow for individual-level stochasticity in predation when analysing data from such experiments, although some have attempted this in a broader setting of population dynamics models [47].

Instead, our approach aims to more directly and explicitly model the searching and consumption of prey. Our model is also stochastic and individual-based to allow for random variation across fish and time in these behaviours. This stochasticity is essential because the number of prey in each treatment of the experiment is relatively few, and only a single fish (the predator) is in each replicate.

We model fish as being in one of two states: **an active preying (AP)** state and a **between-preying (BP)** state. The AP state represents periods when fish are actively seeking prey, while the BP state encompasses times when they are not—for instance, due to digestion, doing something unrelated to preying, or a lack of motivation to feed (in the functional response literature, this is typically classed as a prey “handling” behaviour but we broaden its definition here to recognise it covers a range of other behaviours). We assume that the duration of the AP state depends on prey density: if there are more prey, it takes less time to find and capture a prey item; and that the duration of the BP state is independent of prey density.

The transitions between the AP and BP states can be described using chemical reaction-like equations:

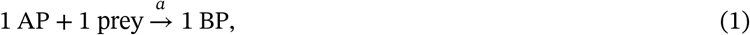

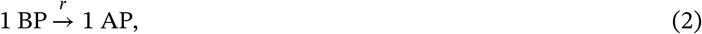

where eq. (1) represents the capturing of a prey item by a fish in the AP state, which leads the fish to transition to the BP state (e.g. when it handles and digests the prey item); the parameter *a >* 0 represents the rate at which this capture occurs; eq. (2) represents the rate at which the fish in the BP state then reverts to the AP state when it then actively seeks prey; the parameter *r >* 0 represents the rate at which this process occurs; *h* = *1/r* represents the average time a fish remains in the BP state, and, in the functional response literature, is known as the *prey handling time*.

For brevity, we refer to *a* as the *rate of prey capture* and *r* as the *rate of prey handling*, whilst recognising that biological interpretation of these rates is wider than this terminology.

Eq. (1) depends on the availability of a prey item, so the rate at which any prey item is captured is given by *Na*, where *N* ≥ 0 is the number of prey still available to capture (i.e. not already eaten). Eq. (2) is *independent* of prey density, so the aggregate rate is just *r* since there is a single predator.

In our experiments, fish are put in enclosures with a given fixed number of initial prey, N_0_ > 0. This means that, initially, the rate of capture of prey is greatest at *aN*_*0*_, which then decreases linearly as prey are consumed. This model, although simple, can produce a range of aggregate outcomes (Fig. 2).

**FIGURE 2.**
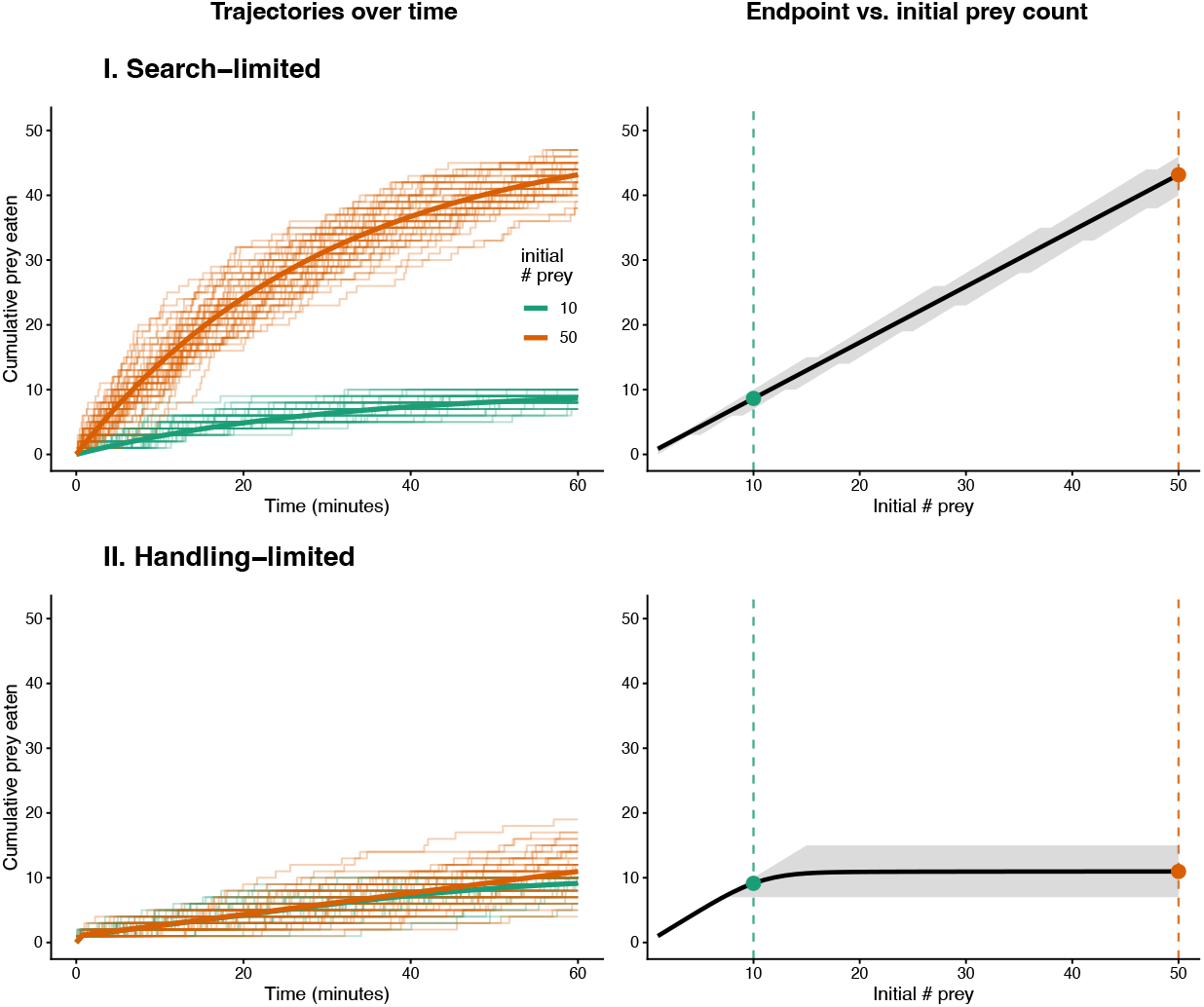
Behaviour of the two-state model in two limiting regimes: a search-limited regime (I), where prey capture is the rate-limiting step, and a handling-limited regime (II), where prey handling limits consumption. Left column: simulated cumulative prey eaten over the one-hour trial (thin lines: individual stochastic realisations; thick lines: mean) for two initial prey densities. Right column: the resulting relationship between the number of prey eaten by the end of the trial and the initial prey density.

#### 2.4.2 Incorporating covariates

The fish are exposed to two conditions: the acclimation conditions (i.e. the pre-experimental conditions) and the experimental conditions (i.e. the conditions the fish experience during the hour when the experiment takes place).

The fish are also heterogeneous; a key source of heterogeneity is their weight, which affects the number of prey fish eat during an experiment—the heavier the fish, the more it eats [48].

The full set of covariates used in our model included fish weight and its squared term, as well as binary variables associated with acclimation and experimental conditions. For each of the acclimation and experimental conditions, four treatment conditions were defined: *ambient, warmed, turbid* and *mixed*. We additionally included a binary variable which captured the *duration* of time the fish were exposed to the acclimation condition (1 week vs. 6 weeks). Interactions between acclimation condition and acclimation duration were also incorporated, as exploratory data visualisations suggested that the magnitude and direction of associations varied with acclimation duration.

We allowed the rates of prey capture, a, and rate of prey handling, r, to vary systematically across replicates according to covariates:

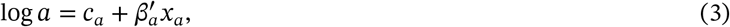

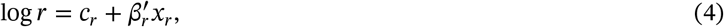

where *x*_*a*_ is a vector of covariates associated with variation in the rate of prey capture and *x*_*r*_ is a vector of covariates associated with variation in the rate of prey handling. By fitting the model to our experimental data, we estimate *c*_*a*_, *β*_*a*_, *c*_*r*_, *β*_*r*_, where *c*_*a*_ and *c*_*r*_ are scalar constants, and *β*_*a*_ and *β*_*r*_ are vectors of coefficients of the same length as *x*_*a*_ and *x*_*r*_, respectively. It was unclear a priori which sets of covariates should be associated with either prey capture (i.e. eq. (3)) or prey handling (i.e. eq. (4)), so we considered two covariate structures, each of which we viewed as equally biologically plausible:

- A model where all covariates except those relating to weight were associated with variations in *a* and only the weight variables were associated with variations in *r*;
- A model where *a* was assumed to be constant (i.e. eq. 3 became just log *a = c*_*a*_) and all covariates were assumed to be associated with variations in *r*.

#### 2.4.3 A subpopulation of voracious foragers

Initial fits of the two-state model systematically under-predicted the fraction of fish that consumed *all* of their available prey. This suggested that, for a substantial share of individuals, prey capture was effectively instantaneous relative to handling, that is, the searching phase contributed negligibly to the outcome. We then extended each model to include a latent subpopulation of *voracious* foragers, corresponding to the limit *a* → ∞ of eq. (1). In this limit a fish is constrained only by prey handling, and the number of prey eaten in a trial follows a shifted, right-truncated Poisson distribution, min(*N*_0_, 1 + Poisson(*r*)) (derived in §C). Each fish belongs to the voracious class with a weight-dependent probability *π*, where logit(*π*) = *γ*_0_ + *γ*_*w*_ *x* weight, and the latent class membership is marginalised out of the likelihood.

We fitted each of the two covariate structures both with and without this voracious-forager component (models with this component we term *mixture* models), giving four candidate models, and chose between them by model comparison (described in the *inference* section).

#### 2.4.4 Inference

In each of our experimental replicates, we record only the number of prey remaining *at the end* of 1 hour. Each experiment had potentially a range of covariates and differing numbers of initial prey, *N*_*0*_. We fitted our models to all these data simultaneously.

We performed Bayesian inference to infer *(c*_*a*_, *β*_*a*_, *c*_*r*_, *β*_*r*_) for each of our models. Bayesian inference requires a prior probability distribution and a likelihood. Rather than adopt nominally uninformative priors, we used weakly informative priors calibrated so that the distribution over prey consumption they induce is consistent with the range observed in the data (a prior predictive check; see Fig. D1). Specifically, we placed Gaussian priors *c*_*a*_ ~ *N*(log 4, 1^2^) and *c*_*r*_ ~ *N*(log 20, 0.5^2^) on the log-rate intercepts, and independent 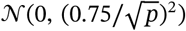 priors on each of the *p* regression coefficients in *β*_*a*_ and *β*_*r*_; continuous covariates (fish weight and its square) were standardised before inference. Each coefficient’s prior standard deviation is set to 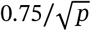 so that the *p* standardised covariates together contribute a fixed total spread (≈ 0.75) to the log-rate regardless of how many there are, keeping the prior-predictive prey counts calibrated and the models comparable; the 0.75 is the target aggregate prior standard deviation of the covariates’ combined contribution to the log-rate, chosen so that at one prior standard deviation they jointly shift the rate by up by a factor of *e*^0.75^ ≈ 2. Priors for the voracious-forager component are given below.

Specifying the likelihood required determining the probability distribution over the number of prey remaining at the end of a given experimental replicate. In §B, we explain how this probability distribution was derived. There was negligible measurement error in the number of prey remaining at the end of each experimental replicate (this simply involved counting the number of bloodworms still present). Uncertainty in the final number of prey remaining then stemmed only from variation inherent in the underlying stochastic reaction equation system.

For the mixture models, we also set priors on the parameters that determined the voracious-feeding-behaviour, *π*. We placed weakly-informative *N*(−1,1.5^2^) and *N*(−0,1.5^2^) priors on *γ*_0_ and *γ*_*w*_; the standard deviation of 1.5 renders the induced prior on the voracious probability gently varying on (0,1) rather than concentrated at the boundaries, while the intercept mean of −1 encodes a weak prior expectation that voracious foragers are a minority (the prior mean for *π ≈* 0.35 and a prior on P(*π* < 0.5) ≈ 0.70).

We also performed prior sensitivity analyses, where we inflated the prior variances and shifted their location. This confirmed that our results were not sensitive to these assumptions (§E).

To estimate the *posterior distribution* over the model parameters, we used Markov chain Monte Carlo (MCMC). For this, we used the differential-evolution adaptive Metropolis (DREAM) algorithm as implemented in PINTS [49], which shares information across a population of chains. To initialise the sampler, we drew a single parameter vector from the prior, optimised it using the CMA-ES algorithm (also implemented in PINTS) to obtain a high-posterior point and started all 40 chains in a tight Gaussian ball (standard deviation 0.3 per coordinate) around that point. Sampling was run in successive segments: the initial optimiser-seeded segment was discarded in full as burn-in and DREAM was continued from its final states (warm-started); we additionally discarded the first 500 iterations of the continued run, retaining 3,500 iterations per chain (140,000 pooled draws). All parameters satisfied 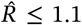, with the largest values (up to ≈1.10) occurring for the attack- and handling-rate intercepts. Tables of parameter estimates with quantiles and 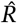 are given in the Appendix. To check the fit of our model to the data, we performed visual *posterior predictive checks*, where we compared the distribution over number of prey remaining (at the end of each experimental replicate) to the modelled equivalent (§D). To perform model comparison, we used an approximate leave-one-out-cross-validation approach as implemented in the loo R package [50].

For discussion of our results, we assess differences between models using a Region of Practical Equivalence (ROPE), which defines a range of values considered too small to be meaningful in practice. Specifically, we examined the posterior distribution of the difference between model estimates and calculated the proportion falling within ±1 minute (for discussions where handling time was the output), our chosen ROPE based on the scale of the analysis. Differences within this interval are interpreted as practically equivalent, while differences outside it indicate a meaningful advantage of one model over the other. This approach allows us to distinguish statistically detectable differences from those that are large enough to matter in practice. We used a 0.95 probability threshold, such that if at least 95% of the posterior mass falls within the ROPE we conclude practical equivalence, whereas if at least 95% falls outside the ROPE on one side we conclude a meaningful difference.

### 2.5 Reproducing the results

The data and code needed to reproduce these results are available on a public GitHub repository: github.com/rccreswell/ functional-response. We facilitate reproducing these results using a bespoke *conda* environment and a data analysis pipeline tool called Snakemake [51].

## 3 Results

Experimental trials were completed for a total of 668 individual fish. Our experimental design produced 224 unique treatment conditions with three fish replicates each. However, for six treatments only two replicates were completed, and for two treatments four replicates were completed. These discrepancies were due to experimenter error.

### 3.1 The model allowing for covariate-associated changes in handling rate with voracious feeding performed best

We compared four models (Table 1) and found that the best-fitting model placed the treatment covariates on the handling rate *r* and included the voracious-forager component. This model was favoured by cross-validation over the alternative covariate structure and the corresponding two-state models without the voracious component (Table 1). Posterior predictive checks for the chosen model (Fig. 3; §D) indicate generally a good correspondence between the observed and simulated data. The only exception was that our model slightly under-predicted the variance in the number of prey consumed (by approximately 16%), most likely reflecting unmodelled variation among individual fish; because the model accurately reproduced mean consumption and the proportions of fish eating all or none of their prey, we expect this to have little influence on the estimated treatment effects, beyond making their credible intervals marginally too narrow.

**TABLE 1.** Model comparison by leave-one-out cross-validation (PSIS-LOO). Each model places the treatment covariates on either the attack rate *a* or the handling rate *r*, with or without a voracious-forager mixture component. “Treatments + weight” denotes the full covariate set on the log rate: acclimation condition, experimental condition, acclimation duration (1 vs. 6 weeks), the acclimation-by-duration interaction, and fish weight (linear and quadratic terms, each variable standardised); “constant” means that rate has an intercept only. 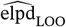 is the expected log-pointwise predictive density (higher is better) and Δelpd is the difference versus the best-fitting model, with its standard error. Rows are ordered best to worst.

| Covariates on $a$ | Covariates on $r$ | Mixture | $\widehat{\text{elpd}}_{\text{LOO}}$ | SE | $\Delta\text{elpd}$ | SE($\Delta$) |
| --- | --- | --- | --- | --- | --- | --- |
| constant | treatments + weight | Yes | −1513 | 45 | – | – |
| treatments + weight | constant | Yes | −1690 | 51 | 177 | 24 |
| treatments + weight | constant | No | −2749 | 217 | 1236 | 203 |
| constant | treatments + weight | No | −2831 | 228 | 1318 | 213 |

**FIGURE 3.**
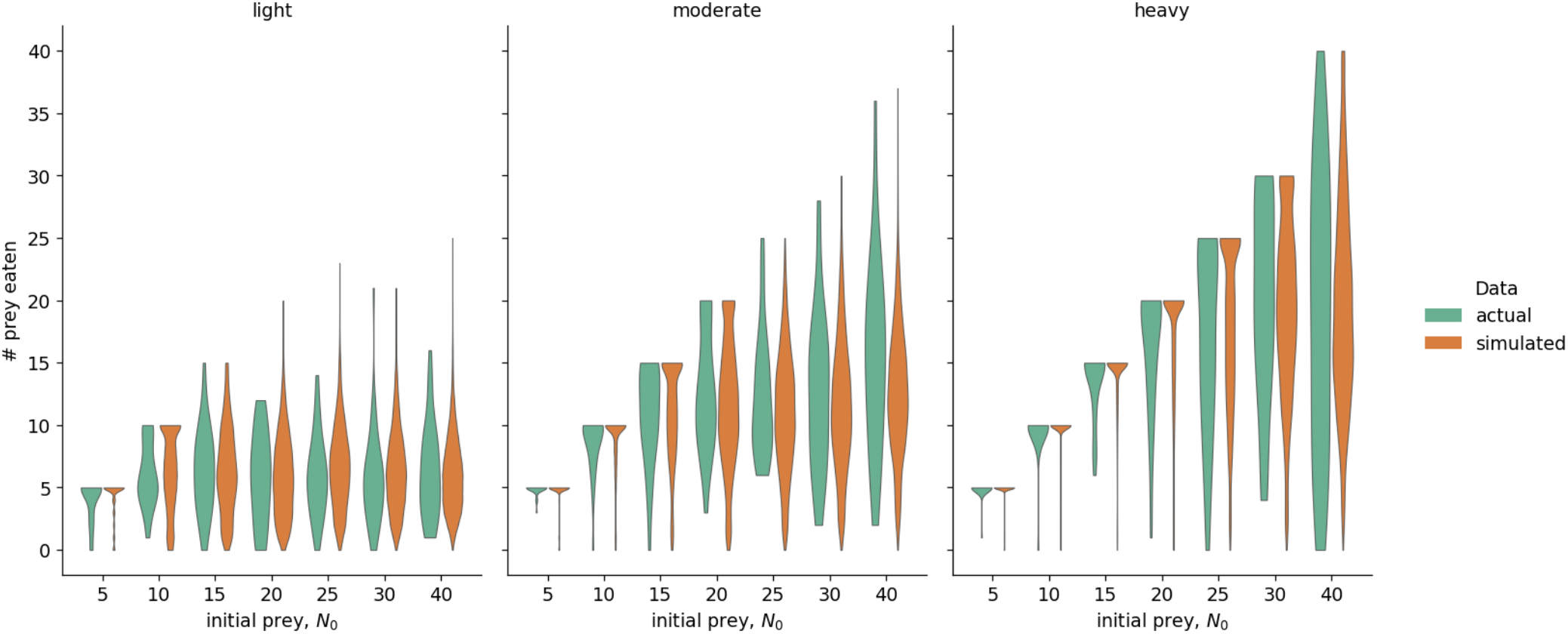
Comparison of actual versus simulated data from the best-fitting model. For each initial prey density *N*_0_, the distribution of the number of prey eaten in the data *(actual*, green) is compared with the distribution simulated from the fitted model *(simulated*, orange), with fish grouped into light, moderate, and heavyweight terciles (i.e. groups of equal size in each of the three categories). A close correspondence between each pair of distributions indicates a good fit.

### 3.2 Prey searching is negligible for most fish

The two-state model on its own substantially under-predicted the number of fish that consumed all of their available prey: across prey densities, 29% of fish cleared their tanks, whereas the two-state model predicted only around 8%. Extending the model to allow a subpopulation of voracious foragers, for whom prey capture is effectively instantaneous (see Methods), resolved this discrepancy (Table D2) and was favoured by cross-validation (Table 1).

The selected model estimated that the great majority of fish, 93% (95% CI: 90–95%), belonged to the voracious class, and this proportion increased with body weight, from approximately 78% for the lightest fish to near 100% for the heaviest (Fig. 4). In other words, for a substantial majority of individuals, the time spent searching for prey was negligible relative to handling time, so that consumption was governed almost entirely by the rate of prey handling, *r*. This provides a quantitative basis for the observation that searching time varied little across treatments: within the confined experimental arenas, prey encounter was so rapid that the searching phase contributed negligibly to feeding for the overwhelming majority of fish.

**FIGURE 4.**
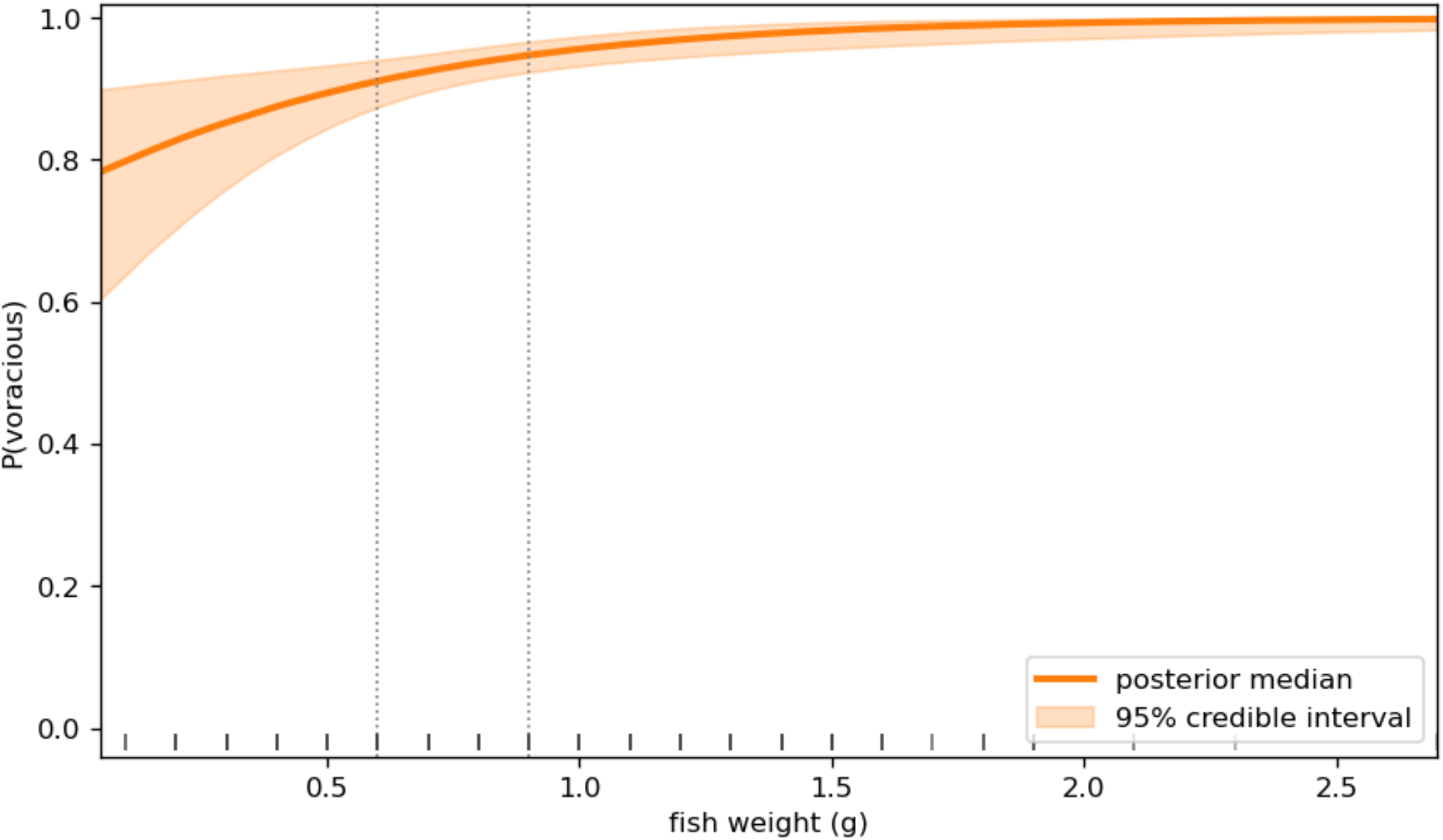
Estimated probability that a fish belongs to the voracious-forager class as a function of bodyweight. The orange line shows the posterior median and the shaded band the central 95% posterior interval. Dotted vertical lines mark the weight tercile boundaries separating the light, moderate and heavy groups.

### 3.3 The effect of experimental treatment on feeding

Controlling for the acclimation condition, we found minimal association between the type of experimental condition and the estimated handling time (Fig. 5A): the corresponding differences in handling time were small and lay within or close to the ±1 minute ROPE (Table 2).

**TABLE 2.** Comparisons of handling-time differences for a fish of median weight. Each cell gives the posterior difference in handling time relative to ambient conditions (median and central 95% credible interval, minutes), evaluated as an *experimental* effect (varying the experimental condition with acclimation held at ambient) and as an *acclimation* effect (varying the acclimation condition with the experiment held at ambient); the final row is the acclimation duration contrast, it has no counterpart in the experimental-condition column because duration was varied only for acclimation. Asterisks give the strength of evidence that the difference lies *outside* the ROPE (a meaningful difference): * P(| Δ | > 1) > 0.95, ** > 0.99, ***> 0.999; no asterisk indicates the difference is not distinguishable from practical equivalence.

| Contrast (vs. ambient) | $\Delta$ handling time (min), median [95% CI] | |
| --- | --- | --- |
|  | Experimental effect | Acclimation effect |
| Warmed, 1 week | +0.5 [+0.1, +1.0] | -1.9 [-2.4, -1.5]*** |
| Warmed, 6 weeks | +0.6 [+0.1, +1.0] | +2.1 [+1.3, +2.9]** |
| Turbid, 1 week | +0.7 [+0.2, +1.2] | -1.7 [-2.2, -1.2]** |
| Turbid, 6 weeks | +0.7 [+0.2, +1.3] | -1.7 [-2.2, -1.2]** |
| Mixed, 1 week | +0.8 [+0.4, +1.3] | -1.8 [-2.3, -1.3]*** |
| Mixed, 6 weeks | +0.9 [+0.4, +1.4] | -0.7 [-1.3, -0.1] |
| Warmed acclimation, 6 vs. 1 week | — | +4.4 [+3.7, +5.2]*** |

**FIGURE 5.**
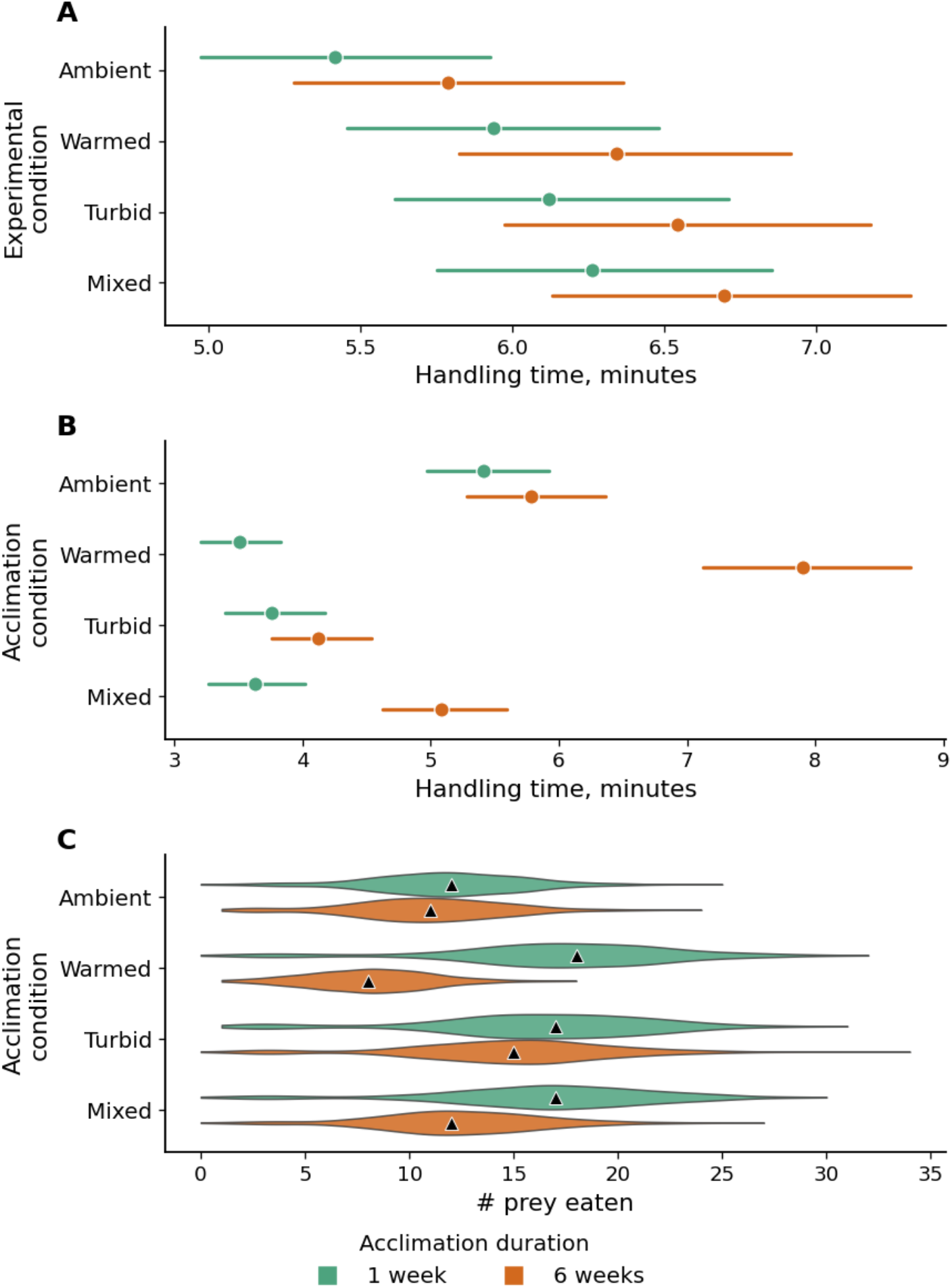
Association between experimental and acclimation conditions and the feeding behaviour of fish. (A) The relationship between the estimated handling time and experimental conditions for fish acclimatised in ambient conditions. (B) The relationship between handling time and acclimation conditions where the experiment was conducted in ambient conditions. (C) The model-predicted number of prey items eaten over the course of a one-hour experiment, where the initial number of prey was assumed to be 40. In all panels, the results are stratified according to the duration of the acclimation period, and the results are generated for a fish of median weight. In A and B, the whiskers show the central 95% posterior intervals for the handling time; the points show the posterior medians for this quantity. In C, triangles show the posterior medians, and the violins provide kernel density estimates of the distributions.

### 3.4 The effect of acclimation treatment and acclimation duration on feeding behaviour

There were substantial differences in the estimated handling time across acclimation treatments and durations (Fig. 5B), with the most pronounced effect under the warmed acclimation treatment. Fish of median weight in the control treatment, maintained under ambient conditions, showed similar handling times regardless of acclimation duration (1 wk: 5.4 min, 95% central posterior interval (95% CI): 5.0–5.9; 6 wks: 5.8 min, 95% CI 5.3–6.4), and consumed comparable numbers of prey (1 wk: 12 prey, 95% CI11–12; 6 wks: 11 prey, 95% CI10-12) (Fig. 5C). In contrast, fish acclimated to warmed water exhibited a strong duration-dependent response: individuals exposed for one week had a handling time of 3.5 minutes (95% CI: 3.2-3.8), whereas those acclimated for six weeks had a substantially longer handling time of 7.9 minutes (95% CI: 7.1-8.7). Relative to ambient conditions at the same acclimation duration, both handling times lay outside our ±1 minute region of practical equivalence (ROPE; Table 2), indicating a meaningful reduction after one week (*P*>0.99)

Acclimation duration and a meaningful increase after six weeks (*P*>0.99); the increase from one to six weeks was itself meaningful (*P*>0.99). Furthermore, our results indicate that fish exposed to warming over longer periods consume considerably more prey than those exposed for shorter periods, and that feeding behaviour differs significantly between both warming treatments and ambient conditions (Fig. 5C).

Fish acclimated to turbid water also showed marked reductions in handling time compared to those in ambient conditions. Consequently, a 1-week acclimation to turbid water reduced handling time by approximately 31% (95% CI 23–38%) for fish of median weight relative to ambient conditions; the reduction after 6 weeks was approximately 29% (95% CI 21–36%); both reductions were meaningful under the ROPE (a meaningful reduction at one week and a meaningful reduction at six weeks, *P>0.99* and *P*>0.99; Fig. 5B).

The effect of combined warming and turbidity (mixed acclimation treatment) was dependent on exposure duration. Fish acclimated under these conditions had shorter handling times than those in ambient conditions, although the magnitude of this effect differed with acclimation length. Fish acclimated for one week exhibited a ~33% reduction in handling time (one-week acclimated handling time: 3.6 min, 95% CI 3.3–4.0), a meaningful difference under the ROPE (a meaningful reduction, *P*>0.99). After six weeks the reduction was only ~12% (six-week acclimated handling time: 5.1 min, 95% CI 4.6–5.6); this difference fell largely within the ±1 minute ROPE, so we found no conclusive difference from ambient conditions at that duration (*P*(|Δ| < 1)=0.83). Fish in the mixed treatment also consumed more prey than those in ambient conditions. Individuals in the one-week treatment consumed more prey on average (17 prey, 95% CI 15–18) than those exposed for six weeks (12 prey, 95% CI 11–3) and both ambient treatments (Fig. 5C). While handling times were broadly similar across different stressor treatments (warming, turbid, and mixed) after short-term exposure, clear differences emerged following acclimation, indicating that longer-term exposure alters the relative effects of these stressors on feeding behaviour.

## 4 Discussion

In this study, we show that the effects of environmental stressors on fish behaviour depend critically on prior exposure conditions. We found that the best predictors for feeding behaviour during the experimental period were not the experimental condition itself, but the acclimation duration and condition the fish were previously exposed to. Across treatments, we found that short-term exposure to warming, turbidity, and their combination, generally increased feeding performance relative to ambient conditions, as indicated by reduced handling times and increased prey consumption. However, these positive effects shifted markedly with prolonged exposure. After six weeks, warming had significant negative effects on feeding performance, whereas turbidity increased feeding rates, albeit to a lesser extent than after short-term exposure. In contrast, the mixed treatment led to intermediate outcomes that matched the feeding performance of fish acclimated in ambient conditions, suggesting that the influences of turbidity and warming partially offset one another. Based on these results, conditions most similar to longer-duration heatwaves are likely to cause the most significant ecosystem changes. Importantly, however, the effects of warming were not simply strengthened with increasing exposure duration, but changed direction: short-term warming increased feeding performance, whereas prolonged warming reduced it. Thus, predictions based on short-term responses alone could substantially misrepresent, or even predict the opposite direction of the ecological effects that emerge with prolonged exposure. A similar importance of heatwave duration has been demonstrated experimentally in sea stars, where prolonged heatwave exposure impaired growth, while shorter heatwaves had relatively limited effects and allowed recovery [52, 53]. In our system, while short heatwaves may initially increase feeding rates and productivity, longer heatwaves could suppress feeding and lead to a subsequent decline in energy intake. As climate change is expected to increase the duration of heatwaves, such shifts from initial ‘boom’ to subsequent ‘bust’ phases may have important consequences for predator-prey interactions and the stability of aquatic food webs.

Sticklebacks occupy a wide range of habitats, including marine, brackish, and freshwater systems, and play an important role in shaping ecosystem dynamics [54]. A growing body of research suggests that, under environmental stress, sticklebacks can increase rapidly in abundance and dominate ecosystems [e.g. 55, 56]. This success is primarily driven by their broad thermal tolerance, rapid reproduction, and generalist feeding strategy [54]. While such resilience may appear beneficial at the species level, it can have substantial ecological consequences. Sticklebacks are mesopredators, and exert top-down control on prey and high densities of stickleback can reduce the abundance of herbivorous zooplankton and fish eggs [56, 57, 58, 59, 60]. This can alter food web structure through multiple pathways: egg predation can reduce recruitment of other fish species [61,62,63,64], while predation on zooplankton can reduce the availability of prey for other predator species leading to resource competition and reduced recruitment success [65, 55].

Across all treatments, we observed a Type II functional response, characterised by an increase in feeding rate at low prey densities, followed by a plateau associated with finite handling time. This type of response implies that predators maintain high consumption rates even at low prey densities, and suggests that there is an increased risk of prey over-exploitation when prey densities are low [25]. Therefore, our results indicate that stressors which decrease handling time and increase predation rate (i.e. 1wk turbidity/warming/mixed and 6wk turbidity) may lead to significant changes in food web structures by reducing the abundance of prey. There is already evidence that such trophic effects can scale up to influence broader ecosystem structure in the Baltic Sea [66]. Conversely, our extended heatwave (6wk warming) scenario may lead to reduced predation rates by sticklebacks and thus more abundant and diverse zooplankton communities [56]. Sustained warming may therefore restructure communities through quite different trophic pathways. Indeed, experimental warming of marine communities has been shown to reduce community biomass and simplify ecological networks through the loss of both trophic and non-trophic interactions [67]. Having more diverse prey could also lead to more larger predators in the ecosystem, so long as those predators do not rely on sticklebacks as their main source of food. However, the reduction in stickleback feeding rate is likely a symptom of decreasing health and fitness. For example, previous studies have shown that sticklebacks kept in warmed water show decreased parental care and reproductive success [68,28], increased disease susceptibility [69, 70, 71], and decreased growth rates [72]. If a species known for its broad thermal tolerance changes its behaviour so drastically under heatwave simulated conditions, it is likely that other species in the same ecosystem will fare similarly or worse as water temperatures approach or exceed their thermal tolerance. Therefore, we believe that the decreased handling time and predation rate associated with the 6wk warmed treatment observed in our study, indicates that these conditions will likely have significant negative impacts on aquatic ecosystems.

Increased predation of zooplankton by sticklebacks can have further significant ecosystem effects. The zooplankton eaten are often key algal grazers and their depletion can lead to increased algal growth and eutrophication, ultimately increasing turbidity [58, 56, 73, 74]. Consequently, the increased prey consumption under turbid conditions observed in our study may contribute to a positive feedback loop, whereby sticklebacks further increase turbidity and eutrophication. This effect may be worsened as stickleback populations grow, given that turbidity has also been shown to increase fecundity in this species [75]. As not all species benefit from turbidity [76], this mechanism may contribute to declines in biodiversity observed in some freshwater systems as stickleback populations expand and displace other species [55].

Crucially, our results reveal that when stressors co-occur, their effects are not simply additive. Rather than amplifying effects in either direction, the combined influence of warming and turbidity in our experiments appears to counteract one another. This is consistent with previous research showing that non-additive interactions between environmental stressors are common, with combined effects frequently being less than expected based on individual effects [77, 78, 79]. Therefore, these conditions may result in more stable feeding dynamics. As human induced environmental change is expected to increase both the duration and intensity of heatwaves and turbidity, this finding challenges the assumption that multiple stressors will necessarily intensify ecosystem degradation. Instead, it raises the possibility that, under certain conditions, their interaction could moderate stickleback-driven changes in food webs and even confer a degree of resilience, particularly for species that respond positively to turbidity.

In this study, we do not directly test the mechanisms underlying changes in feeding behaviour associated with environmental stressors, but we can speculate. The reduction in handling time observed after one week of warming is likely driven by increases in metabolic rate. Previous studies have shown that higher temperatures can accelerate metabolic rate in sticklebacks, resulting in greater feeding and growth rates [80]. But fish have an upper thermal limit beyond which metabolic performance declines [72], that may account for the longer handling time observed after prolonged exposure. The mechanisms underpinning behavioural responses to turbidity are less well understood. The lack of a behavioural change in fish moved from clear acclimation conditions to turbid experimental conditions (Fig. 5A) suggests that turbidity does not strongly disrupt foraging behaviour in sticklebacks. This contrasts with findings in other species, such as the coral reef triggerfish *(Rhinecanthus aculeatus)*, where turbidity has been shown to immediately reduce foraging efficiency, even in the presence of other sensory cues (e.g. olfaction) [81, 76]. Sticklebacks are known to be visual predators, but from a sensory ecology perspective, our results suggest that turbidity may not limit prey detection, despite it impacting other visually mediated behaviours such as courtship activity, mate choice, and shoaling [34, 35, 82]. Several mechanisms may explain why feeding performance was enhanced under turbid conditions. First, the prey used in our experiments (bloodworms) are relatively conspicuous and presented within a confined space. These conditions may minimize the sensory challenge of prey detection. In addition, we used bentonite clay as a source of turbidity. This material has a slightly red colour and while the increased turbidity will have reduced visual contrast, it may have enhanced or maintained the saliency of the prey colour cue [83]. An alternative mechanism is that turbidity reduced the visual detection distance of prey, but that this effect was offset through behavioural compensation. Increased turbidity has been shown to reduce predator vigilance in the three-spined stickleback [84], allowing individuals to increase prey search effort. A final option is that sticklebacks may adjust sensory sensitivity in response to turbidity over time. For example, they could switch to a potentially more reliable sensory cue in low visibility conditions such as olfaction. However, experiments suggest that olfaction is used primarily to detect the presence of prey rather than locate it, and may instead act to increase the motivation to forage [43].

An additional contribution of this study is the modelling framework itself. Despite its relatively simple structure, the model performed well across the range of experimental conditions and was able to detect biologically meaningful differences in feeding behaviour from inherently noisy behavioural data. This is particularly valuable given the model was fit only to endpoint observations (number of prey eaten after an hour) rather than the timing of feeding events occurring within trials. We did not use the camera footage to produce within-experiment feeding data owing to there being insufficient feeding events within the segments filmed. Thus, the model represents prey consumption as a simple Markovian process, with two memoryless states corresponding roughly to prey searching and prey processing. In reality, these behaviours are likely to depend on idiosyncratic mechanisms such as an individual’s past experience and physiology, details which cannot be extracted from the data collected in this study. This means that our model infers the within-experiment feeding patterns rather than being informed by it, and future such studies might consider these richer data for informing such models. However, despite its simplicity, across the various checks we performed and reported, our model provided a reasonable explanation of the observed patterns.

Finally, the results of this study highlight an important implication for behavioural research. We found that behavioural change was not driven by the immediate experimental conditions, but instead by the acclimation conditions experienced prior to testing. This suggests that short-term experimental approaches, such as functional response assays, may in effect be capturing prior environmental history rather than responses to the experimental treatment itself. This underscores the importance of accounting for the environmental and experiential history of test subjects when interpreting results [85]. Moreover, the persistence of altered behaviour after fish were removed from their acclimation conditions indicates that prolonged exposure to stressors can have lasting effects, even once conditions return to baseline. In the context of future environmental change, where stressors such as heatwaves and rapid-onset turbidity events are expected to become more frequent, this suggests that behavioural impacts may persist beyond the duration of the disturbance, with recovery occurring over extended timescales.

## Author Contributions

**Cait Newport:** Conceptualization (equal); Investigation (equal); Methodology (equal); Resources (lead); Software (supporting); Supervision (lead); Visualisation (equal); Writing - Original Draft Preparation (lead). **Ellen Morley:** Investigation (equal); Methodology (equal). **Richard Creswell:** Software (supporting); Writing - Review & Editing. **Michelle Jackson:** Conceptualization (equal); Methodology (supporting); Resources (supporting); Writing - Review & Editing (equal). **Ben Lambert:** Methodology (equal); Software (lead); Visualisation (equal); Writing - Review & Editing (equal).

## Acknowledgments

We thank Mike M. Webster for helpful advice on stickleback populations and husbandry in the U.K. and Tim Coulson for reading an early draft of this manuscript.

## Funding

This research received support through Schmidt Sciences, LLC (to C.N.) and the Natural Environment Research Council (NERC; NE/V001396/1 to M.J.)

## Conflicts of Interest

The authors declare no conflicts of interest.

## Data Availability Statement

The data and code supporting the findings of this study are available on GitHub at github.com/rccreswell/ functional-response. An archived version of the data and code will be made available in Zenodo prior to publication.

## Notes

### Competing Interest Statement

The authors have declared no competing interest.

### Summary of Updates

Correction to the spelling of the name of author Richard Creswell.

